# A conserved maternal-specific repressive domain in Zelda revealed by Cas9-mediated mutagenesis in *Drosophila melanogaster*

**DOI:** 10.1101/210187

**Authors:** Danielle C. Hamm, Elizabeth D. Larson, Markus Nevil, Kelsey E. Marshall, Eliana R. Bondra, Melissa M Harrison

## Abstract

In nearly all metazoans, the earliest stages of development are controlled by maternally deposited mRNAs and proteins. The zygotic genome becomes transcriptionally active hours after fertilization. Transcriptional activation during this maternal-to-zygotic transition (MZT) is tightly coordinated with the degradation of maternally provided mRNAs. In *Drosophila melanogaster,* the transcription factor Zelda plays an essential role in widespread activation of the zygotic genome. While Zelda expression is required both maternally and zygotically, the mechanisms by which it functions to remodel the embryonic genome and prepare the embryo for development remain unclear. Using Cas9-mediated genome editing to generate targeted mutations in the endogenous *zelda* locus, we determined the functional relevance of protein domains conserved amongst Zelda orthologs. We showed that neither a conserved N-terminal zinc finger nor an acidic patch were required for activity. Similarly, a previously identified splice isoform of *zelda* is dispensable for viability. By contrast, we identified a highly conserved zinc-finger domain that is essential for the maternal, but not zygotic functions of Zelda. Animals homozygous for mutations in this domain survived to adulthood, but embryos inheriting these loss-off-function alleles from their mothers died late in embryogenesis. These mutations did not interfere with the capacity of Zelda to activate transcription. Unexpectedly, these mutations generated a hyperactive form of the protein and enhanced Zelda-dependent gene expression. These data have defined a protein domain critical for controlling Zelda activity during the MZT, but dispensable for its roles later in development, for the first time separating the maternal and zygotic requirements for Zelda. This demonstrates that highly regulated levels of Zelda activity are exclusively required for establishing the developmental program during the MZT. We propose that tightly regulated gene expression is essential to navigate the MZT and that failure to precisely execute this developmental program leads to embryonic lethality.

**AUTHOR SUMMARY:** Following fertilization, the one-celled zygote must be rapidly reprogrammed to enable the development of new, unique organism. During these initial stages of development there is little or no transcription of the zygotic genome, and maternally deposited products control this process. Among the essential maternal products are mRNAs that encode transcription factors required for preparing the zygotic genome for transcriptional activation. This ensures that there is a precisely coordinated hand-off from maternal to zygotic control. In *Drosophila melanogaster,* the transcription factor Zelda is essential for activating the zygotic genome and coupling this activation to the degradation of the maternally deposited products. Nonetheless, the mechanism by which Zelda functions remains unclear. Here we used Cas9-mediated genome engineering to determine the functional requirements for highly conserved domains within Zelda. We identified a domain required specifically for Zelda’s role in reprogramming the early embryonic genome, but not essential for its functions later in development. Surprisingly, this domain restricts the ability to Zelda to activate transcription. These data demonstrate that Zelda activity is tightly regulated, and we propose that precise regulation of both the timing and levels of genome activation is required for the embryo to successfully transition from maternal to zygotic control.

## INTRODUCTION

During the first hours following fertilization, the zygotic genome is transcriptionally silent, and maternally deposited products control early development. These maternal products establish regulatory networks that enable the rapid and efficient transition from two specified germ cells to a population of totipotent cells, which give rise to a new organism. This dramatic change in cell fate is coordinated with the transition from maternal to zygotic control of development, resulting in a complete reorganization of the transcriptome of the embryo. The maternal-to-zygotic transition (MZT) is comprised of two essential and coordinated events, (I) transcriptional activation of the zygotic genome, and (II) destabilization and degradation of maternally supplied RNAs [1–4]. The concerted action of two RNA clearance pathways ensures the timely elimination of maternally deposited transcripts [5–11]. The first is a maternally encoded pathway that initiates the degradation of maternal RNAs in the absence of fertilization and zygotic transcription. The second pathway is zygotically triggered and contributes to maternal RNA clearance near the end of the MZT. Thus, transcriptional activation of the zygotic genome is precisely coordinated with degradation of the maternally provided products [5,10,12]. Regulation of these events is required for development, as failure to undergo this transition is lethal to the embryo. Nonetheless, the mechanisms that precisely control the timing and levels of gene expression necessary to successfully navigate this dramatic developmental transition remain to be elucidated.

In *Drosophila melanogaster,* the MZT occurs over the first few hours of development. The transcription factor Zelda (ZLD; Zinc-finger early *Drosophila* activator) is a critical regulator of the MZT, and its absence is lethal to the embryo [13–17]. *zld* transcripts are maternally deposited and robustly translated following fertilization leading to ubiquitous protein expression in the pre-blastoderm embryo [14,17,18]. ZLD binds to thousands of *cis*-regulatory modules and is required for transcriptional activation of the zygotic genome [13–15]. ZLD is necessary for gene expression both early and late during the MZT; ZLD drives expression of a small number of genes as early as the eighth mitotic division and is required for the later activation of hundreds of genes during the major wave of zygotic genome activation at mitotic cycle 14 [13]. Among the genes that require ZLD for expression are components of the RNA degradation pathways that destabilize maternal RNAs [14,16]. These ZLD-target genes include several zygotically expressed miRNAs and lncRNAs, including the miR-309 cluster of miRNAs that mediates degradation of over one hundred maternally loaded RNAs [16,19]. Thus, maternally supplied *zld* is essential for zygotic genome activation and maternal mRNA decay, driving the coordinated transition from maternal-to-zygotic control. ZLD is also required zygotically, such that embryos homozygous for a deletion in *zld* die late in embryogenesis [14,17].

Maternally deposited *zld* encodes a protein of 1596 amino acids, including six C2H2 (Cys-Cys-His-His motif) zinc fingers, but no known catalytic activity (Fig. 1) [14,17,20]. In tissue culture, ZLD is a robust transcriptional activator, and this function requires the C-terminal cluster of four zinc fingers that comprise the DNA-binding domain and a low-complexity region proximal to this domain [20]. Functional data combined with phylogenetic analysis supports a shared role for ZLD in genome activation among insects and crustaceans [20–25]. Thus, we were surprised to discover that while transcriptional activation is a conserved function of ZLD, in cell culture this activity does not require highly conserved regions in the N-terminus, including two of the C2H2 zinc fingers and an acidic patch [20,25]. A truncated splice isoform of *zld* is also conserved throughout the *Drosophila* genus. This variant is expressed in late embryos and in larvae, but lacks coding sequence for three of the four C-terminal zinc fingers in the DNA-binding domain and is therefore unable to bind DNA (Fig 1A) [20,26–28]. The conservation of these additional domains and splice isoforms suggests a potential function that has been retained through evolution, but which may not have been evident in cell culture.

**Figure 1.**
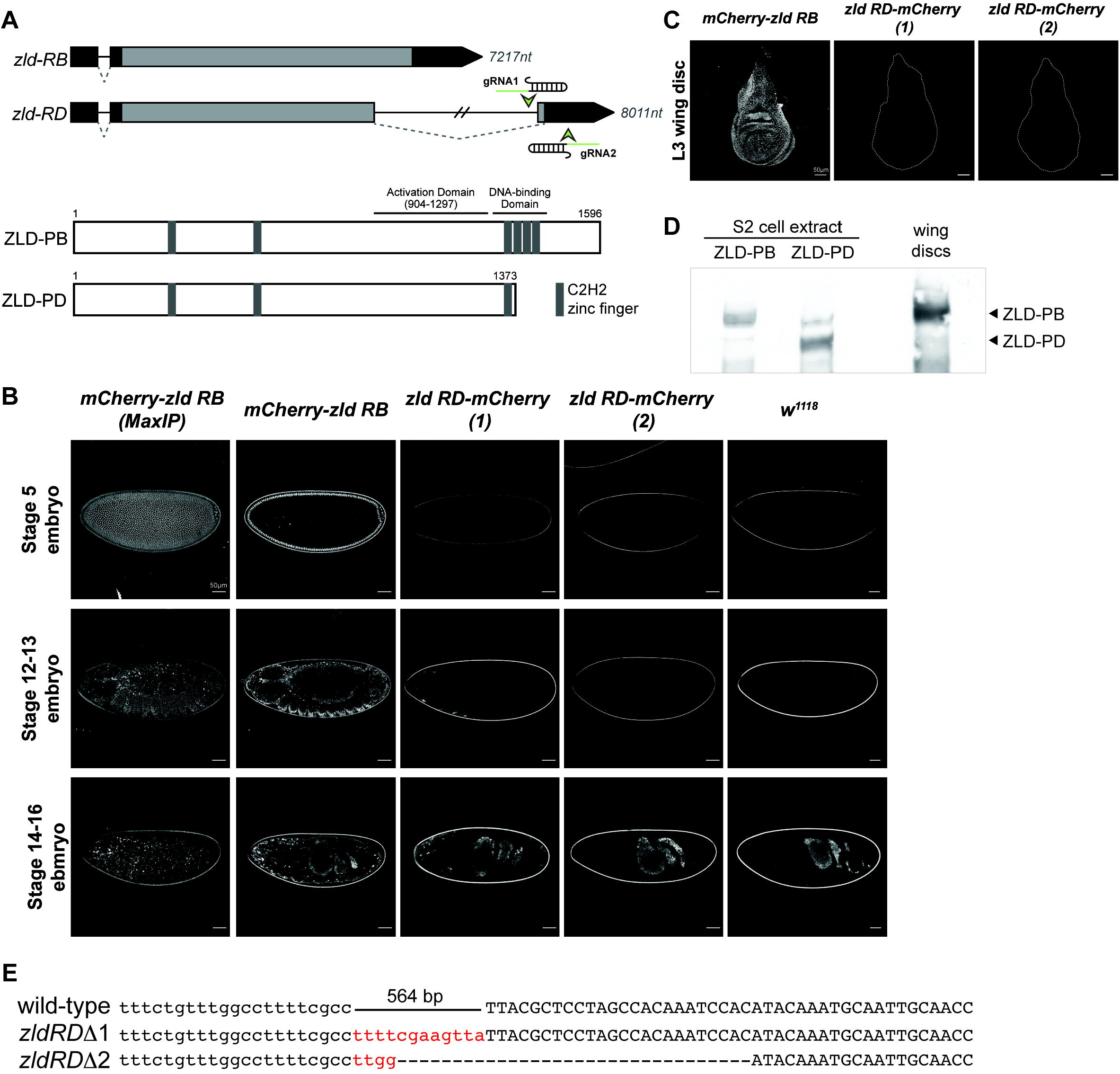
ZLD-PB is the dominant isoform expressed both in the embryo and imaginal disc. (A) Schematic of predicted transcripts from the *zld* locus (*above*). Boxes indicate exons with coding sequence (light gray) and untranslated regions (dark gray). The length of the resulting mRNA is given in nucleotides (nt). Two gRNAs flanking the downstream exon of *zld-RD* used to generate an isoform specific deletion are shown. Schematics of the predicted protein products for each splice variant (*below*) with amino acid numbers in *D. melanogaster* ZLD shown above. Above are the approximate locations of the transcriptional-activation and DNA-binding domains as demonstrated in Hamm et al (2015) [20]. (B) Confocal images of embryos homozygous for either an N-terminal mCherry-tagged ZLD or for ZLD-PD-mCherry at stages 5, 12-13, and 14-16. The first column shows the maximum projection images for mCherry-ZLD expressing embryos. All other images show a single confocal slice. ZLD-PD-mCherry (1) and (2) indicate embryos from two distinct editing events. (C) Confocal images of mCherry-ZLD isoforms demonstrating the endogenous ZLD and ZLD-PD specific expression in third instar larval wing discs. Outlines show the borders of the wing discs as determined by transmitted light images. All images shown are the maximum projection. (D) Immunoblot for ZLD on S2 extract from cells expressing either ZLD-PB or ZLD-PD or total lysate from third instar wing discs. (E) Sequence of *zld-RD* following Cas9-mutagenesis demonstrating removal of the splice acceptor and coding sequence. Small insertions (red sequence) and deletions (dashed lines) shown. Lower case letters indicate intron sequence. Capital letters indicate exonic sequence.

Our recent development of techniques for Cas9-mediated genome engineering in *Drosophila* enabled us to directly test the roles of these conserved features of ZLD *in vivo* [29]. Previous approaches to investigate the *in vivo* function of specific protein domains relied largely upon the use of transgenes, which do not always adequately reflect the endogenous expression patterns, levels, or alternative splice isoforms. We therefore developed a rapid and efficient means to screen for Cas9-mediated point mutations. Generation of specific point mutations allowed us to interrogate the function of conserved features of *zld in vivo.* Using a combination of epitope tags and targeted deletions, we demonstrated a truncated *zld* isoform was unlikely to be translated and was not required for viability in *D. melanogaster*, despite being conserved amongst Drosophilidae. We generated targeted loss-of-function alleles for conserved domains in the N-terminus, including the two zinc fingers and the acidic patch. The first C2H2 zinc finger (ZnF1) and the acidic patch (EDD) were dispensable for viability. To our surprise, the second zinc finger (ZnF2) was required for maternal, but not zygotic function of ZLD. Embryos laid by mothers homozygous for mutations in the second zinc finger died late in embryogenesis. Contrary to our expectations, mutations in this zinc finger resulted in a hyperactive version of ZLD that caused precocious activation of the zygotic genome and increased degradation of maternal transcripts. Together these data demonstrate, for the first time, a separable function for maternally and zygotically expressed ZLD and suggest that the early embryo is exquisitely sensitive to ZLD activity such that too little or too much activity results in embryonic lethality.

## RESULTS

### The 1596 amino acid ZLD protein is the principal isoform expressed throughout development

*zld* transcripts are present throughout the *Drosophila* life cycle. They are strongly expressed during oogenesis, resulting in ubiquitous protein expression in the pre-blastoderm embryo. Subsequently, *zld* is zygotically expressed in the developing embryonic germ layers, nervous system, imaginal disc primordia and in larval wing and eye discs [17,27,28,30]. In the pre-blastoderm embryo *zld* contains a single, unspliced open reading frame and a single five-prime intron (Fig 1A). This open reading frame translates into the 1596 amino acid protein product ZLD-PB. In addition to this maternally deposited isoform, a truncated isoform, *zld-RD*, which contains a second unique downstream exon and alternative splice junction, is expressed during zygotic development [26–28]. This alternatively spliced isoform codes for a 1373 amino acid protein lacking three of the four C-terminal zinc finger motifs required for DNA binding (Fig 1A) [20,31]. The truncated product resulting from translation of the *zld-RD* isoform acts as a dominant negative when co-expressed with the 1596 amino acid isoform in cell culture [20]. Nonetheless, it was unknown whether this shorter isoform was translated to form a protein product *in vivo* and if so, whether it was expressed in the same cell as the longer 1596 amino acid isoform, which would be required for any dominant negative effect on ZLD activity.

To determine the expression pattern of a protein product from the *zld-RD* isoform, we used Cas9-mediated genome engineering to tag each of the two protein isoforms with mCherry. Because we had previously shown that the N-terminal 900 amino acids of ZLD are dispensable for activating transcription in cell culture [20], we tagged the N-terminus to avoid interfering with protein function. Flies carrying this mCherry tag are homozygous viable and fertile demonstrating that the tag does not interfere with any of the essential functions of ZLD. Since all *zld* splice isoforms encode proteins with a shared N-terminus, expression of the N-terminal mCherry tagged protein is indicative of the expression pattern of all known ZLD isoforms. To specifically determine the expression pattern of a protein product of the shorter *zld-RD* isoform, we engineered an mCherry tag upstream of the stop codon in the downstream exon that is specific to *zld-RD* (Fig 1A). This addition did not affect the *zld-PB* isoform. Like flies carrying the N-terminal fluorescent tag, these flies were also homozygous viable and vertile.

We imaged stage 5, 12-13, and 14-16 embryos homozygous for either the N-terminal mCherry tag or the *zld-RD* specific mCherry tag to determine the expression patterns of ZLD protein products (Fig 1B). *zld-RB* is ubiquitously expressed in the pre-blastoderm embryo, while post-blastoderm expression is limited to the tracheal primordium, central nervous system (CNS), and midline neurons [14,27]. Similar to the expression pattern for the mRNA, the N-terminally tagged protein was expressed throughout embryogenesis (Fig 1B). We did not detect fluorophore expression from either of the two strains containing the mCherry-tagged *zld-RD* (Fig 1B). Thus, despite high levels of *zld-RD* in the CNS of stage 12-16 embryos [27], the ZLD-PD isoform does not appear to be expressed. We detected gut auto fluorescence in all genotypes, including control *w*^*1118*^ embryos.

To investigate additional tissues that might express ZLD-PD at later stages of development, we imaged imaginal wing discs from third instar larvae (L3). Previous reports had suggested that ZLD-PD was expressed in larval tissues [28]. Using our engineered fly lines, we could detect ZLD expression in several L3 tissues, including ubiquitous, nuclear expression in imaginal wing discs (Fig 1C). By contrast, we could not detect mCherry expression in the lines specifically tagging ZLD-PD (Fig 1C). In addition to *zld-RD,* a second splice isoform of *zld, zld-RF,* has been identified and is predicted to produce a protein product very similar to the predicted product of *zld-RD*. The evidence for *zld-RF* is weaker than for *zld-RD,* and it has been speculated to be the result of a cloning artifact [26,27,32]. Nonetheless, both truncated isoforms have been reported to be expressed in the wing imaginal disc [28]. To determine if any truncated protein product is translated from either the *zld-RD* or *zld-RF* isoforms, we immunoblotted protein extract from wing imaginal discs using our antibody that recognizes all isoforms of ZLD [20]. We identified only a single isoform, corresponding to the 1596 amino acid protein (Fig 1D). This evidence suggests that neither *zld-RD* nor *zld-RF* isoforms are translated to a stable protein in the wing imaginal disc.

The *zld-RD* splice isoform is conserved throughout the *Drosophila* genus [26,27], suggesting a retained function. Thus, it remained possible that the truncated RNA or the splicing reaction was instrumental to *zld* function and could explain the conservation, even if the protein isn’t stably expressed. To test this possibility, we determined the *in vivo* effects of eliminating the *zld-RD* isoform by using Cas9-mediated mutagenesis to delete the splice acceptor and downstream coding region of *zld-RD* (Fig 1E). We obtained two strains carrying the deleted sequence, both of which were viable and fertile (Fig 1E). Because the exons encoding *zld-RF* are within the required longer isoform, we were unable to make a deletion targeting only this isoform as we did for *zld-RD*. Therefore, we cannot rule out the possibility that this isoform is important *in vivo*. While *zld-RD* is expressed in multiple post-blastoderm tissues as an RNA [26–28], our data demonstrated that this truncated splice-isoform is not essential for development and is not abundantly translated.

### Cas9-mediated genome engineering enables *in vivo* functional analysis of conserved protein domains

Because the 1596 amino acid ZLD-PB isoform is the predominantly expressed form of ZLD, we investigated the functional requirements of domains within this large transcription factor. ZLD-PB is comprised of six C_2_H_2_ zinc fingers and many low-complexity regions, but no identifiable enzymatic domains. Alignment of ZLD orthologs showed sequence conservation within insects of all six zinc fingers as well as an N-terminal acidic patch (Fig 2A–C) [20,25]. We previously demonstrated that the cluster of four C-terminal zinc fingers constituted the DNA-binding domain, and the low-complexity domain just N-terminal to the DNA-binding domain mediated transcriptional activation (Fig 1A) [20]. Therefore, the regions required for both DNA binding and transcriptional activation were encompassed within the 600 C-terminal amino acids of ZLD [20], while the functional significance of the conserved N-terminal zinc-fingers and acidic domain was unknown. Because domains under high evolutionary constraint often possess important structural or functional roles, we hypothesized that these highly conserved domains might have an essential developmental function that was missed in our previous cell-culture assays.

**Figure 2.**
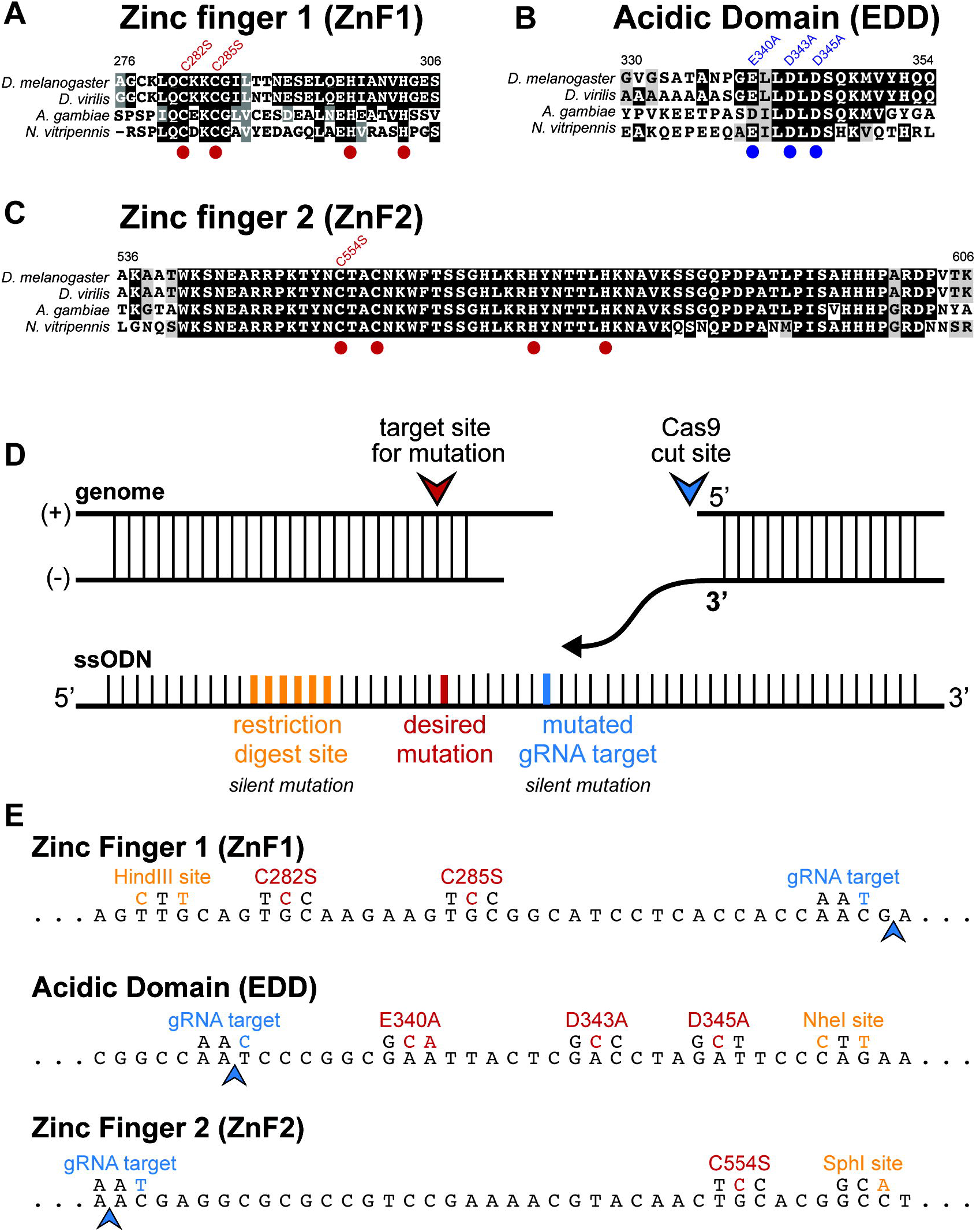
Cas-9 mediated mutation of highly conserved domains in ZLD. (A-C) Alignment of amino acid sequences of ZLD protein domains from four insect species (*Drosophila melanogaster, Drosophila virilis, Anopholes gambiae, Nasonia vitripennis*). Red dots indicate conserved cysteine and histidine residues in the C_2_H_2_ zinc finger domains. Blue dots indicate the conserved aspartate and glutamate residues in the acidic domain. Point mutations generated in the endogenous *zld* locus by Cas9-mediated genome engineering are shown above. Numbers above indicate amino acids in *D. melanogaster* ZLD. (D) Schematic of the experimental design for homology directed repair from a single-stranded donor oligonucleotide (ssODN) following a Cas9-mediated, double-strand break. Silent restriction site (yellow), silent mutation in the gRNA target site (blue) and the desired point mutation (red) are shown. (E) Wild-type DNA sequence coding for the targeted ZLD domains (*below*) with mutated nucleotides generated by homology directed repair (*above*) are shown. Arrowheads indicate Cas9 cleavage site.

The recent development of Cas9-mediated genome editing has allowed us to facilely create point mutants *in vivo*. This strategy enabled efficient creation of endogenous mutant alleles to probe the functional importance of individual protein domains of interest. A single-stranded donor oligonucleotide (ssODN) and a single gRNA construct were injected into flies expressing Cas9 to create loss-of-function mutations in the highly conserved N-terminal domains. We developed a streamlined protocol to molecularly screen for the desired mutations; ssODNs contained both the desired coding mutations and silent mutations that generated a restriction digest site not found in the endogenous locus, allowing for screening by PCR and restriction enzyme digest (Fig 2D,E). Instead of creating deletions, we introduced point mutations in the conserved N-terminal domains with the purpose of maintaining overall protein stability. To disrupt the zinc-finger domains, we mutated a subset of the zinc-chelating residues in each of the N-terminal zinc fingers. Within the acidic domain, we mutated the conserved glutamate and aspartate residues to alanine to abrogate the negative charge in the domain. Using this streamlined strategy, we generated three distinct mutant alleles, individually disrupting each of these conserved protein domains and allowing us to interrogate protein structure and function *in vivo* (Fig 2E).

### The highly conserved second zinc-finger in ZLD is essential to support embryonic development

We assessed the viability and fertility of each of the mutants we generated by counting the number of homozygous flies carrying the mutations as compared to heterozygous siblings (Fig 3). Flies homozygous for mutation of either the first zinc finger (ZnF1) or the acidic domain (EDD) were viable to near wild-type levels (Fig 3A). Both homozygous males and females were fertile. Similarly, hemizygous males carrying mutations in zinc finger 2 (ZnF2) are viable, albeit to a reduced degree, and were fertile. Homozygous ZnF2 mutant females were viable at reduced levels, but, in contrast to their male counterparts, were sterile (Fig 3A). Homozygous *zld*^*ZnF2*^ females lay fertilized embryos that arrest late in embryogenesis during stage 17 after tracheal branches have clearly formed. A subset of males and females carrying only *zld*^*ZnF2*^ had small, malformed eyes, suggesting additional developmental process were affected by the mutation. Thus, contrary to our expectations loss-of-function mutations in all three regions were dispensable for viability even though their conservation suggested they were required for ZLD function. Our mutational analysis delineated discrete maternal and zygotic functions for ZLD; maternal deposition of *zld* mutant for ZnF2 was lethal to the embryo, causing arrest late in embryogenesis, whereas zygotic expression of *zld* with a disruption in ZnF2 supported development to adulthood.

**Figure 3.**
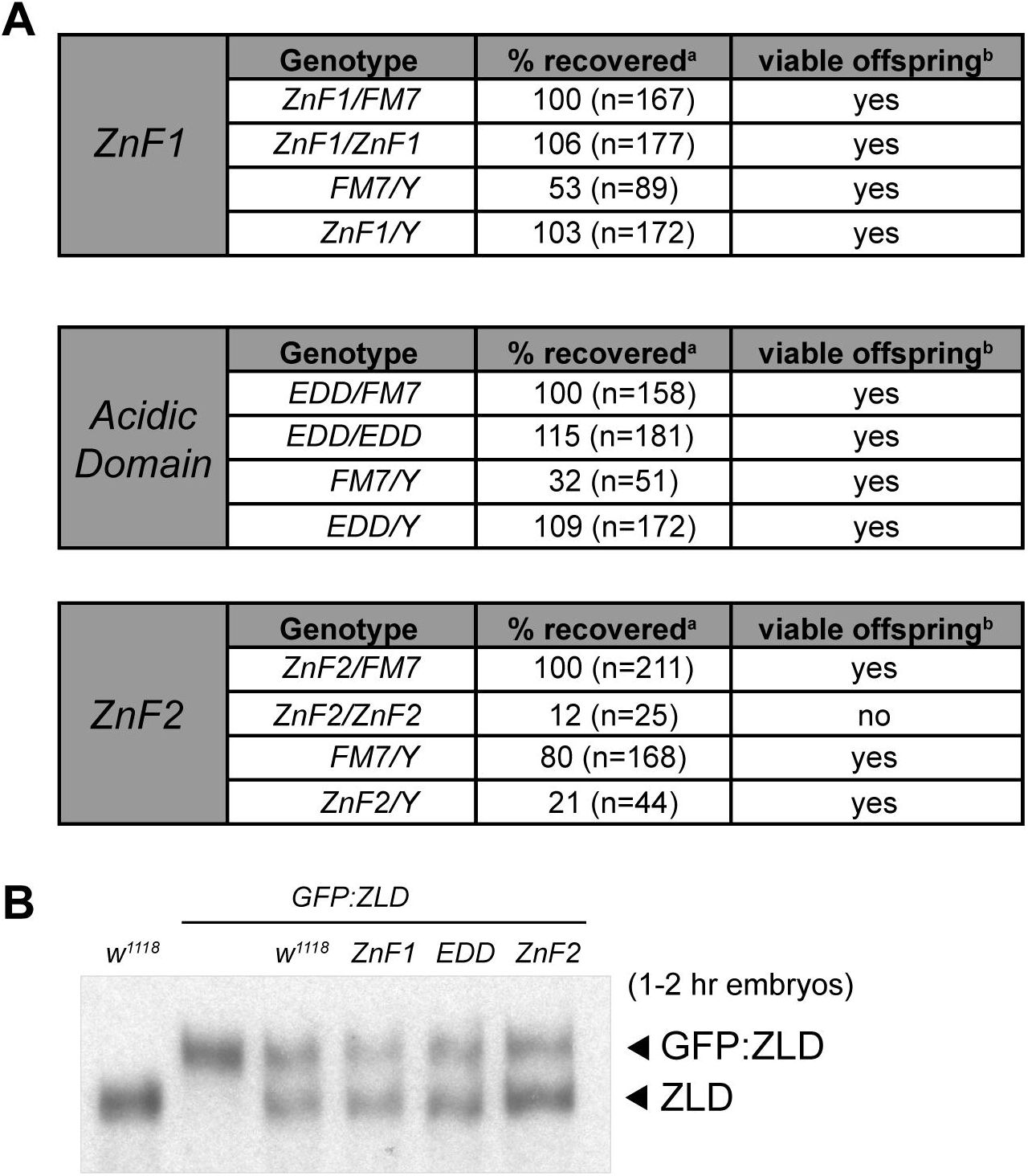
The second zinc finger in ZLD is required for maternal, but not zygotic function. (A) Viability and fertility of heterozygous and homozygous animals carrying point mutations in zinc finger 1 (ZnF1), the acidic domain (EDD), or zinc finger 2 (ZnF2). ^a^Percent recovered was calculated in comparison to *FM7/+* siblings. Percent based on Mendelian inheritance. ^b^Viablity was determined by crossing to a *w*^*1118*^ mate. (B) Immunoblots with anti-ZLD antibodies demonstrate wild-type levels of protein expression from alleles harboring point mutations in conserved domains. Protein levels were assayed on total protein lysate from embryos 1 – 2 hours after egg laying (AEL) maternally inheriting both the mutated allele and a GFP-tagged allele that is fully functional, allowing internal normalization.

To determine if protein stability was disrupted in any of the mutants, we compared expression of each mutant allele to a GFP-tagged endogenous, wild-type allele in a heterozygous embryo. Embryos were laid by heterozygous females such that they received maternal deposition of RNA encoding both ZLD protein variants. Equivalent amounts of protein were expressed from alleles carrying mutations in either ZnF1, ZnF2, or the acidic domain as compared to the GFP-tagged control (Fig 3B). Furthermore, the observed phenotype for the ZnF2 mutant was retained in a trans-heterozygote carrying a deletion in *zld* (*zld*^*294*^) (S1 Table). Thus, the maternal-effect lethality associated with maternal inheritance of *zld* mutant for ZnF2 was not a result of protein instability or a background mutation, but instead was the result of changes in ZLD function.

### Mutations in zinc finger 2 create a novel hyperactive allele

To test the functional importance of these conserved domains on ZLD-mediated transcriptional activation, we used our previously established cell-culture system to trainsiently express ZLD mutants and assay their ability to activate luciferase reporters [20]. ZLD with mutations within either ZnF1 or the acidic domain was able to activate transcription to a level similar to the wild-type protein (Fig 4), consistent with these mutations producing viable and fertile flies (Fig 3A, S1 Table). The single amino acid change (C554S) in the ZnF2 mutant significantly hyperactivated the *scute* reporter, resulting in luciferase activity almost 4-fold greater than wild type. None of the ZLD proteins activated gene expression from a mutant promoter, confirming the specificity of the assay. Immunoblots confirmed the expression of all mutated proteins was at approximately equivalent levels (Fig 4). These data suggest that ZnF2 may serve as an inhibitory domain that regulates the level of ZLD-mediated transcriptional activation. This conclusion is further supported by our previous data demonstrating that truncations to the N-terminus of ZLD that remove ZnF2 elevated transcriptional output [20].

**Figure 4.**
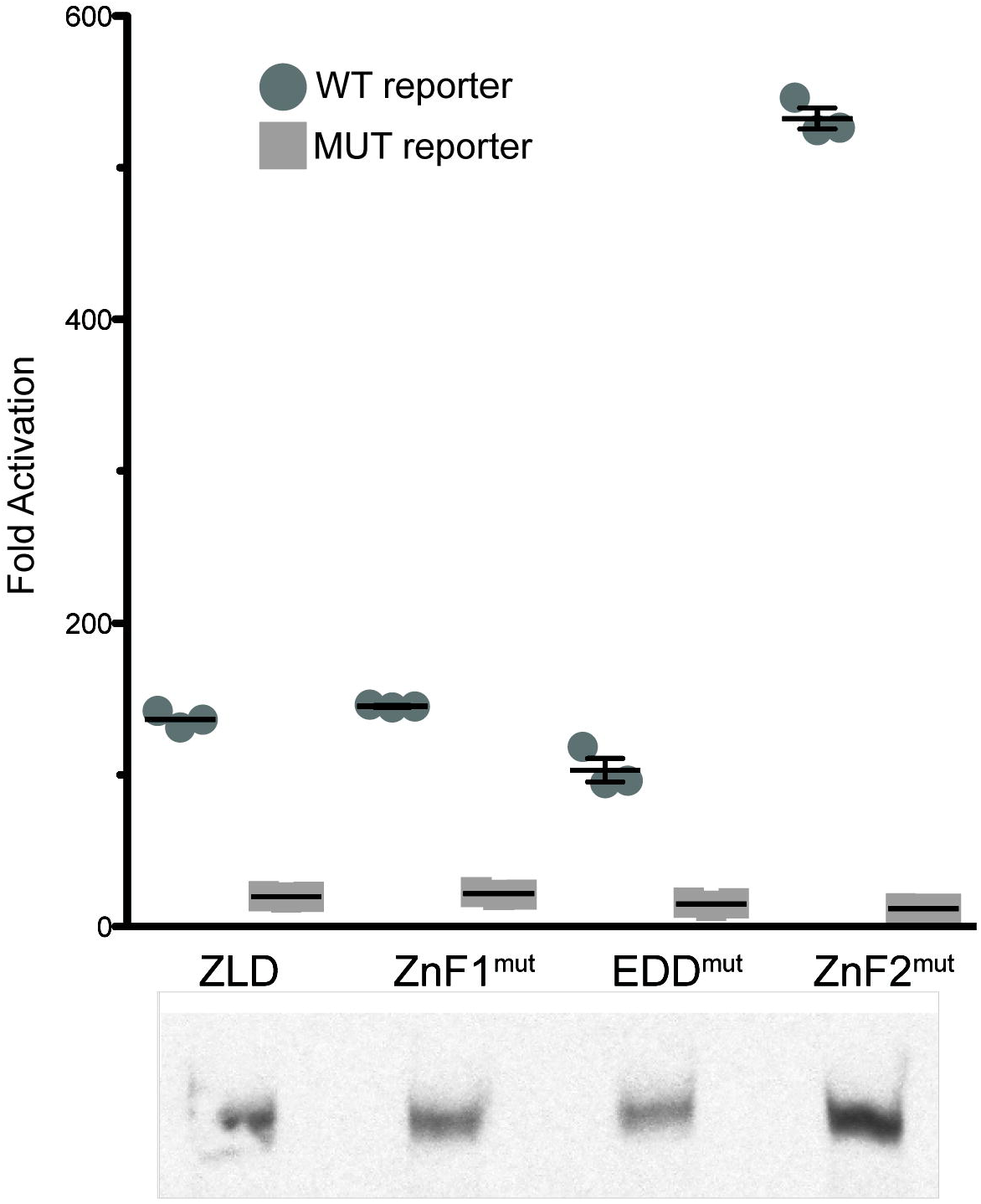
Mutation in the second C_2_H_2_ zinc finger of ZLD results in increased ability to activate transcription. Fold activation of luciferase reporters driven by either a wild-type *scute* promoter (WT) or one with mutations in the ZLD-binding sites (MUT). Immunoblot with anti-ZLD antibodies demonstrates comparable levels of ZLD expression. n = 3, error bars indicate +/− standard deviation.

Prior studies demonstrated that either overexpression of maternal *zld* or the loss of maternally deposited *zld* resulted in defects in nuclear division in the blastoderm embryo [17]. The similarity of the loss-of-function and overexpression phenotypes suggests that the early embryo is sensitive to the precise levels of ZLD activity and that both too little and too much activity is detrimental to embryonic development. To confirm the impact of ZLD overexpression, we used *mat-*α*-GAL4* to drive overexpression of a *UASp-zld* transgene. Overexpression of maternally deposited *zld* caused a late embryonic lethal phenotype similar to that of animals inheriting maternal *zld*^*ZnF2*^, albeit at a lower frequency (S1 Fig). Based on our tissue culture data and the fact that *zld* overexpression phenocopies the ZnF2 mutation, we propose that disruption of zinc finger 2 hyperactivated ZLD protein and that this increased activity was lethal to the embryo.

### Conserved residues outside of the C_2_H_2_ zinc finger are required for regulating ZLD activity

The second zinc finger in ZLD is the most highly conserved domain in the entire 1596 amino acid protein [25]. To determine whether conserved residues outside of the zinc-chelating amino acids of zinc finger 2 were essential for function, we mutated four conserved residues (F561, S563, Y571 and N578) to alanine, generating the *zld*^*JAZ*^ allele (Fig 5 A-C). These residues were chosen because they are shared between ZLD ZnF2 and the consensus sequence for JAZ (Just Another Zinc finger)-domains (pfam: zf-C2H2_JAZ), a domain initially identified in the mammalian double-stranded RNA-binding zinc finger protein JAZ (Fig 5A) [33]. Alanine substitutions in the JAZ domain did not affect protein stability (Fig 5D). Animals homozygous for *zld*^*JAZ*^ were viable at reduced frequencies, and males were fertile (Fig 5E). Females homozygous for this allele were sterile, laying embryos that later died (Fig 5E), phenocopying the *zld*^*ZnF2*^ mutants (Fig 3A, S1 Table). Cell-culture assays further demonstrated that mutating the JAZ-like domain hyperactivated transcription, similar to the serine substitution in C554 (Fig 5F). Thus, both zinc-chelation and residues conserved within the JAZ zinc finger domain are critical for negatively regulating the ability of ZLD to activate transcription.

**Figure 5.**
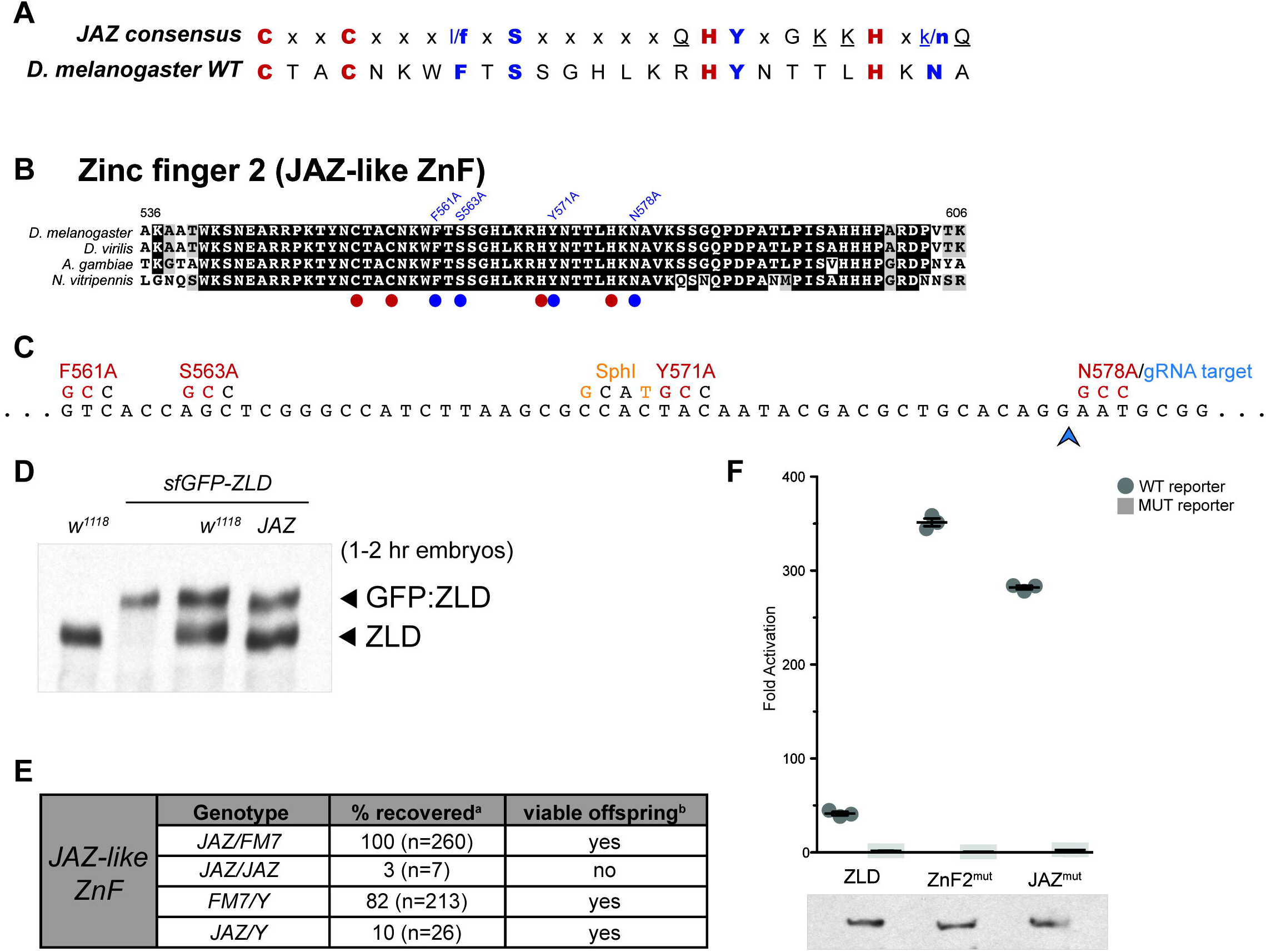
Conserved residues in the second zinc finger domain shared with JAZ-like zinc fingers are essential for wild-type ZLD activity. (A) Alignment of the second zinc finger of ZLD with the consensus amino acids in JAZ domains. Bold indicates amino acids shared between the consensus and ZLD with red indicating the zinc-chelating cysteines and histidines and blue indicating additional shared residues. Underlined residues are those that contact double-stranded RNA. (B) Alignment of amino acid sequence of second zinc finger in ZLD with other insect species. Red dots indicate conserved cysteine and histidine domains in the C_2_H_2_ zinc finger. Blue dots indicate the amino acids conserved in JAZ zinc fingers. Point mutations generated in the endogenous *zld* locus by Cas9-mediated genome engineering are shown above. (C) Wild-type DNA sequence coding for the targeted ZLD domains (*below*) with mutated nucleotides generated by homology directed repair (*above*) are shown. Blue arrowhead indicates Cas9 cleavage site. (D) Immunoblot with anti-ZLD antibodies demonstrates wild-type levels of protein expression from the allele harboring point mutations in the conserved JAZ-domain amino acids. Protein levels were assayed on total protein lysate from embryos 1 – 2 hours after egg laying (AEL) maternally inheriting both the mutated allele and a GFP-tagged allele that is fully functional, allowing internal normalization. (E) Viability and fertility of heterozygous and homozygous animals carrying point mutations in the JAZ domain amino acids. ^a^Percent recovered was calculated in comparison to *FM7/+* siblings. Percent based on Mendelian inheritance. ^b^Viablity was determined by crossing to a *w*^*1118*^ mate. (F) Fold activation of luciferase reporters driven by either a wild-type *scute* promoter (WT) or one with mutations in the ZLD-binding sites (MUT). Immunoblot with anti-ZLD antibodies demonstrates comparable levels of ZLD expression. n = 3, error bars indicated +/− standard deviation.

### ZnF2 is essential for regulating maternal mRNA clearance and zygotic genome activation during the MZT

The cell-culture assays demonstrated that the JAZ-like ZnF2 negatively regulated ZLD activity. This raised the possibility that the lethality in embryos inheriting *zld* with mutations in this domain might result from hyperactivation of ZLD targets. To determine the functional consequences of the *zld*^*ZnF2*^ allele on early gene expression, we performed mRNA-sequencing on hand-sorted stage 5 embryos with wild-type maternal *zld* (*w^1118^*) or *zld*^*ZnF2*^ (C554S). The high degree of reproducibility amongst the three replicates (S2 Fig) allowed us to identify genes misexpressed in embryos inheriting the mutated version of *zld* (Fig 6). We identified 287 genes that were up-regulated in the *zld*^*ZnF2*^ mutant and 270 genes that were down-regulated (Fig 6A and S2 Table). Stage 5 embryos possess both mRNAs that have been deposited by the mother along with newly transcribed zygotic mRNAs. To distinguish between these two classes of mRNAs, we used previously published data to determine whether the mis-regulated genes were maternally, zygotically or both maternally deposited and zygotically expressed (mat-zyg) [34]. 74% (n = 212) of up-regulated genes were zygotically expressed, including those expressed exclusively in the zygote and those maternally deposited and zygotically expressed (Fig 6B). By contrast, only 11.5% (n = 31) of the genes that were down-regulated were zygotically expressed. 73% (n = 197) of the down-regulated genes were maternally deposited (Fig 6B). Thus, the majority of the up-regulated genes were zygotically expressed while the majority of the down-regulated genes were maternally deposited.

**Figure 6.**
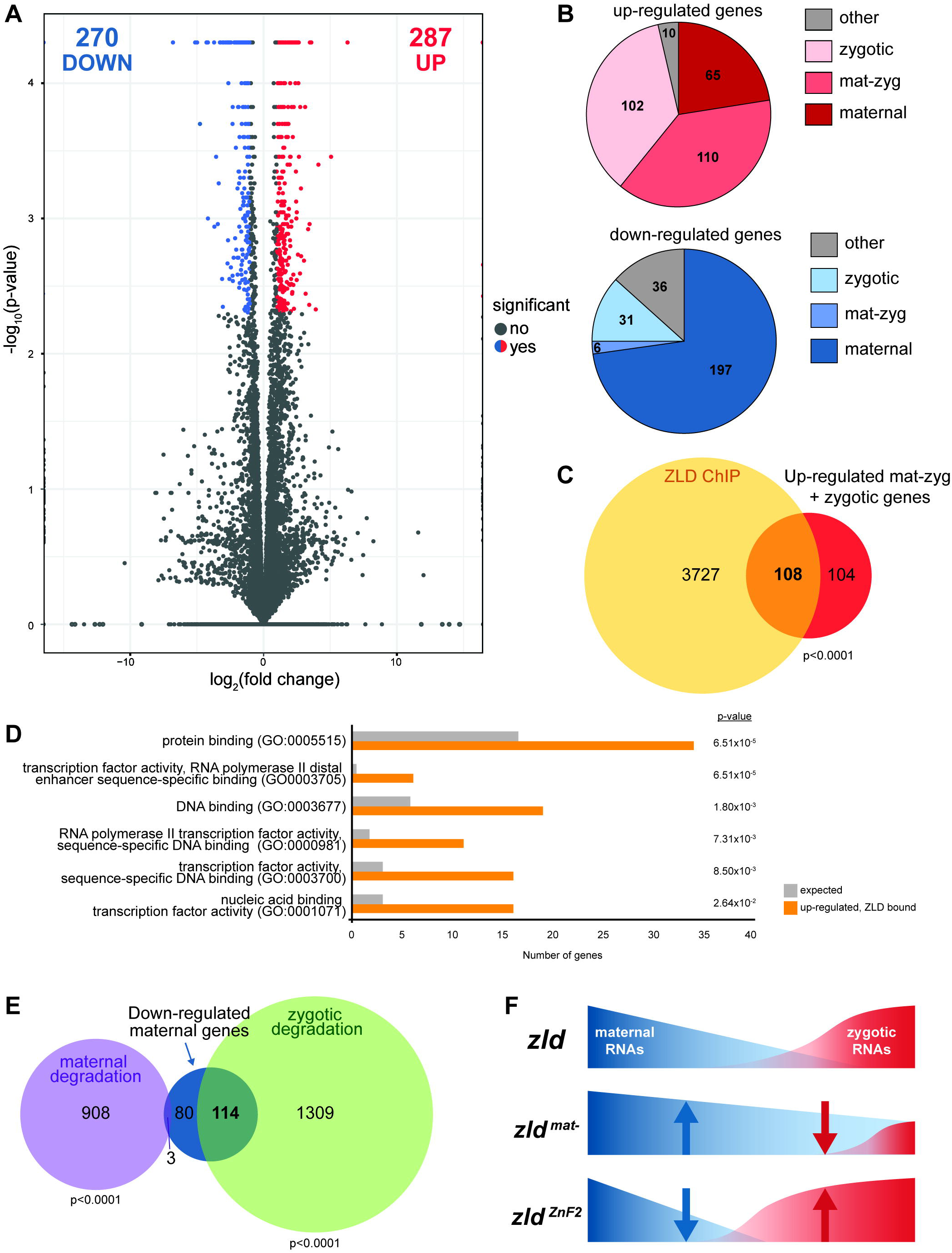
Mutation of the second zinc finger results in both increased zygotic genome activation and maternal RNA degradation during the MZT. (A) Volcano plot showing log2 fold change values (x-axis) by –log10 corrected p-values (y-axis) for all genes identified by RNA-seq. Data represent a comparison of gene expression in embryos laid by either mothers with C554S mutation in zinc finger 2 (*zld*^*ZnF2*^) or by *w*^*1118*^ mothers. Genes that are significantly altered in expression (p-value < 0.05, > 2 fold change) are indicated as red dots (up-regulated) or blue dots (down-regulated). (B) Numbers of up-regulated (red) and down-regulated genes that are maternally expressed (maternal), zygotically expressed (zygotic) or both maternally and zygotically expressed (mat-zyg). Classification of gene expression is as defined in Lott et al. 2011 [34]. (C) Venn diagram showing the overlap of up-regulated, zygotically expressed genes and genes bound by ZLD (as defined by ChIP-seq in Harrison et al. 2011 [13]). (D) GO term enrichment for up-regulated genes associated with ZLD-binding sites. (E) Venn diagram showing the overlap of down-regulated, maternally expressed genes with the sets of genes subject to maternal or zygotic degradation pathways as defined in Thomsen et al. 2010 [10]. p-values are calculated by Fisher’s exact test. (F) Model of the effects on maternal and zygotic gene expression over the MZT due to maternal inheritance of *zld* mutants.

ZLD is required for transcriptional activation of hundreds of zygotic genes during early embryogenesis [13–15,35]. Thus, we tested whether the hyperactive *zld*^*ZnF2*^ allele up-regulated expression of direct ZLD-target genes. We used our previous ZLD ChIP-seq data to identify ZLD-bound regions in the stage 5 embryo and associated them with the nearest gene to identify 3836 potential direct ZLD targets [13]. More than half of the genes up-regulated in embryos inheriting the *zld*^*ZnF2*^ allele overlapped with likely direct ZLD targets (p < 0.0001), suggesting that these genes were directly hyperactivated by the mutant ZLD protein (Fig 6C). To determine the regulatory networks influenced by ZLD^ZnF2^ hyperactivity, we identified enriched Gene Ontology (GO) terms for the 108 likely direct targets. The most enriched GO terms were related to transcription-factor activity, DNA binding, and RNA Pol II activity (Fig 6D). Misexpression of these genes may therefore affect multiple downstream processes required for embryonic development, ultimately leading to the late-stage lethality of these embryos.

Zygotic gene activation is coordinated with the degradation of maternally deposited RNAs during the MZT. Two sets of machinery remove maternally deposited transcripts from the early embryo with one functioning just after fertilization and one functioning later during genome activation [2,5]. The early decay pathway is encoded by maternal factors and is triggered by egg activation. The late-decay pathway is encoded by zygotic factors expressed at the onset of zygotic genome activation [10,12]. Because we found an enrichment for maternally deposited mRNAs amongst the down-regulated transcripts in the embryos inheriting maternal *zld*^*ZnF2*^, we hypothesized that mRNAs in these mutant embryos might be precociously degraded due to hyperactivation of the zygotic genome. To test this, we determined whether the down-regulated mRNAs corresponded to genes subject to either the early or late decay pathways [10,12]. 58% (n = 114) of the down-regulated maternal mRNAs overlap with mRNAs subject to degradation late during the MZT while just 1.5% (n = 3) were degraded early in the MZT (Fig 6E). These data support a model whereby ZLD^ZnF2^ hyperactivates a set of zygotic genes and that this leads to precocious decay of a set of maternally deposited mRNAs (Fig 6F).

## DISCUSSION

The dramatic changes in cell fate that occur during the MZT require precise coordination of activation of the zygotic genome and degradation of the maternally deposited products that drive the initial stages of embryogenesis. For development to proceed this transition must be smoothly executed. In *Drosophila*, ZLD is required for this transition and facilitates activation of hundreds of zygotic genes. Using a combination of evolutionary analysis, Cas9-mediated genome editing and high-throughput sequencing, we identified an essential regulatory domain within ZLD. Our data demonstrate a maternal-specific function for the highly conserved second zinc finger and suggest that the early embryo is exquisitely sensitive to precise regulation of ZLD activity.

### Structure-function analysis of ZLD using targeted Cas9-mediated mutagenesis

*D. melanogaster* have been a premier organism for studies of gene regulation and development for over a century, but studies have been limited by the inability to precisely engineer mutations in the genome using homologous recombination. Our establishment of Cas9-mediated genome engineering in *D. melanogaster* overcame this limitation [29]. Here we have used this facile method of gene editing to identify the functional domains of the essential transcription factor ZLD. We developed a molecular screening strategy that enabled us to generate four distinct mutations to directly query the necessity of conserved protein domains. Editing the endogenous locus provided confidence that any phenotypes we observed were not due to differences in levels or localization of gene expression that might result from the use of a transgene. This was supported by confirmation that all the mutations we generated were expressed at endogenous levels. Thus, we are confident that the absence or presence of a clear mutant phenotype represented the endogenous requirement for a specific protein domain. Our use of genome editing to determine the requirements for specific protein domains within ZLD highlights more generally the power of Cas9-mediated editing to characterize protein structure and function. The easy PCR-based screening approach described here allows for the generation and identification of novel alleles in as little as one months time, providing an additional powerful tool to study gene function in *Drosophila*.

Cas9-mediated genome editing also enabled us to specifically determine the protein expression pattern from a conserved splice isoform of *zld* that is predicted to produce a truncated protein product. Using a combination of an isoform-specific mCherry tag, a targeted deletion, and immunoblot, we clearly demonstrated that while *zld-RD* may be expressed as an RNA it is not translated at detectable levels in either the embryo or the larval wing disc and is not required for viability (Fig 1). Visualization of an N-terminal mCherry tag that marks all possible ZLD isoforms demonstrated that ZLD remains expressed in embryos well past the MZT (Fig 1 B-D). Thus, the 1596 ZLD-PB isoform that binds DNA and drives transcriptional activation is likely the predominant protein product at all stages of development and in all cell types.

### The JAZ-like zinc finger domain regulates ZLD transcriptional activity

A single *zld* ortholog with a set of highly conserved domains is found within the genomes of insects and some crustaceans. These ZLD orthologs are required for embryonic development and transcriptional activation within multiple insect species [20,25,36]. We had previously shown that ZLD-mediated transcriptional activation in *Drosophila* cell culture did not require either of the conserved N-terminal C_2_H_2_ zinc fingers or a recently identified conserved acidic patch [20,25]. Here, we used Cas9-mediated genome editing to test the functional significance of these conserved domains *in vivo* by generating point mutations that were likely to result in loss of function. We individually mutated both conserved N-terminal zinc fingers as well as the acidic patch (Fig 2). Because coordination of zinc ions plays an essential structural role in zinc finger domains [37], the cysteine-to-serine mutations are likely to lead to structural changes that abrogate function. Similarly, removing acidic residues from the acidic patch is likely to disrupt any interactions that relied on the negative charge of these residues. For example, it has been suggested that this negatively charged domain might contact positively charged histones [25], and the alanine substitutions would be expected to block this interaction. Mutation of either the first zinc finger or the acidic patch did not disrupt the ability of ZLD to activate transcription in culture (Fig 4), consistent with our previous cell-culture studies [20], and mutant flies homozygous for these mutations were viable and fertile (Fig 3). These domains are therefore not essential for ZLD-mediated transcriptional activation in *D. melanogaster*. However, given the conservation of these domains, they likely had an ancestral function that might be retained in other insects or an as of yet unidentified activity in *D. melanogaster*. In distantly related species, *Tribolium castaneum* and *Rhodnius prolixus,* knockdown of *zld* after the MZT resulted in extensive morphological defects. These defects included impaired segment generation from the growth zone, a structure unique to these short-germ band insects and not shared in long-germ band insects, such as *Drosophila* [25]. Thus, it is possible that there is an ancestral function of ZLD that is retained in short-germ band insects and requires these conserved domains.

We demonstrated that the second zinc finger of ZLD is required for female fertility. This zinc finger is the most highly conserved domain of the entire protein and has similarity to the double-stranded RNA-binding, JAZ-like zinc finger family [17,20,25,33]. We generated two distinct loss-of-function alleles by mutating either a required zinc-chelating cysteine or four residues that are shared with the JAZ zinc finger domains. These mutations convey a maternal-effect lethality due to an increase in ZLD-mediated transcriptional activation (Fig 4 – 6). Thus, we propose that this domain suppresses the ability of ZLD to activate transcription. While ZnF2 has the canonical architecture of the JAZ-like C_2_H_2_ zinc finger, it lacks a positively charged lysine residues that is conserved in double-strand RNA-binding zinc fingers and is thought to be required for RNA binding [38]. Therefore, it is unlikely that this domain functions through interaction with double-stranded RNA. Nonetheless, this domain likely interacts with a protein partner, a nucleic acid, or intramolecularly within ZLD to effect its suppression of transcriptional activation. To date, no such interactions within ZLD or between ZLD and a protein or RNA partner have been identified.

### ZnF2 serves as an inhibitory domain essential for regulating gene expression during the maternal-to-zygotic transition

*zld* is required as a maternally deposited mRNA that is translated following fertilization. Embryos lacking maternally deposited *zld* die early in embryogenesis due to a failure to undergo the MZT [14]. Embryos homozygous mutant for zygotic *zld* die late in embryogenesis [14,17], but the cause of this lethality is currently unknown. While ZLD is required throughout embryogenesis for viability, it was previously unclear if there were functions distinctly required at either stage of development. The maternal-effect lethality we demonstrate for mutations in the second zinc finger provides the first evidence of separable functional requirements for maternal and zygotic ZLD. Using both tissue culture assays and RNA-sequencing, we showed that this highly conserved zinc finger suppresses the ability of ZLD to activate transcription. Thus, early embryonic development is particularly affected by these mutations. Based on these data, we propose that the early embryo is exquisitely sensitive to ZLD activity such that too little or too much is lethal to the embryo.

One possible explanation for the maternal-specific nature of mutations in ZnF2 is that a cofactor that binds this domain and suppresses activity is only expressed in the early embryo. In this case, loss-of-function alleles would lead to loss of cofactor binding specifically in the early embryo. The fact that we identified increased ZLD-mediated transcriptional activation by ZnF2 mutants in S2 cells demonstrates that any such cofactor must also be expressed in these cells. Because S2 cells are derived from late-stage embryos and *zld* is not normally expressed in these cells [39], we propose that any factor that mediates the ZnF2-mediated inhibition is likely broadly expressed during development. Rather than stage-specific expression of a cofactor, we suggest that the maternal-specific nature of the allele is the result of distinct features of the early embryo that makes this time in development particularly sensitive to ZLD activity. Supporting this hypothesis, we have previously demonstrated that ZLD protein levels are controlled in the early embryo and that ZLD levels increase when its activity is first required [13]. Furthermore, in all insects studied only a single *zld* ortholog has been identified despite extensive expansion of other transcription factor families [25]. This suggests that even a single extra copy of the *zld* locus may be detrimental to development.

Both the mutations we generated in zinc finger 2 and the overexpression of maternally deposited *zld* result in late embryonic lethality rather than lethality during the MZT when the resulting protein is expressed. Our RNA-sequencing data suggest that mutation in ZnF2 results in precocious activation of the zygotic genome (Fig 6). This is supported by our demonstration that not only are a subset of ZLD-target genes up-regulated in these mutants, but that there is a coordinated decrease in levels of maternally deposited mRNAs. These maternal mRNAs are enriched for those that depend on zygotic transcription for degradation. Early activation of ZLD-target genes and precocious degradation of maternal mRNAs leads to misregulation of genes essential for subsequent patterning of the embryo, including transcription factors as evidenced by our GO term analysis. These changes in gene expression timing and levels are likely to have cascading effects that result in the subsequent lethality later in embryonic development.

The first few hours of embryonic development require the rapid transition from a specified germ cell to a population of totipotent cells. A maternally provided program controls the transition from maternal to zygotic control to drive these dramatic changes in cell fate. The coordination of this event is facilitated by the requirement for zygotically encoded proteins to degrade a population of maternally provided transcripts. As a master regulator of transcriptional activation in the zygote, ZLD is essential for executing this transition with precision. Here we have shown that ZLD activity is strictly controlled through a conserved, maternal-specific inhibitory domain – the lack of ZLD, its overexpression, or its hyperactivity via mutations in the inhibitory domain are all lethal to the embryo. We propose that rapid transitions in cell fate, such as those that occur during the MZT, must be precisely executed and that this requires tight control of both the levels and activities of the master regulators of these cell fate changes.

## MATERIALS AND METHODS

### Antibodies and plasmids

Antibody used for immunoblots were rabbit anti-ZLD antibodies at 1:750 [40]. Firefly luciferase reporters containing the *scute* promoter were previously described in Hamm et al. 2015 [20]. *Actin:renilla* was used to control for transfection efficiency [41]. Protein coding regions were cloned into pAc5.1 (Invitrogen) for protein expression in S2 tissue culture cells. PCR was used to amplify the open reading frames of *zld* containing point mutations from genomic DNA obtained from engineered CRISPR mutants. Amplified products were cloned into pAc5.1 for expression in *Drosophila* S2 cells.

### Cell culture and dual-luciferase assays

*Drosophila* S2 cells were cultured at 25°C in Schneider’s Media (Life Technologies) supplemented with 10% Fetal Bovine Serum (Omega Scientific) and antibiotic/antimycotic (Life Technologies). Transfections were performed in triplicate in 24 well dishes with a total of 300 ng of DNA using Effectene Transfection Reagent (Qiagen). Luciferase assays were performed using the Dual Luciferase assay system (Promega). Fold activation was determined by comparison with transfections using a plasmid containing the *actin* promoter but no expression sequence. Representative data sets are shown with error bars indicating the standard deviation.

### Fly stocks

Fly strains used in this study include: *w*^*1118*^, *vasa-Cas9* (BDSC #51324*); w*^*1118*^*; Cyo, P{Tub-PBac\T}2/wg*^*Sp-1*^ (BDSC #8285), *mat-*α*-GAL4-VP16* (BDSC #7062), *wzld*^*294*^*/FM7* [14]*. UASp-zld* flies were made by PhiC31 integrase-mediated transgenesis into the M{3xP3-RFP.attP}ZH-86Fb docking site (BDSC #24749). The following *zld* mutant alleles were generated using Cas9-mediated genome engineering (outlined in detail below): *sfGFP-zld*, *mCherry-zld-RB*, *zld-RD-mCherry*, *zld-RD* deletion, *zld*^*ZnF1*^ *zld*^*EDD*^, *zld*^*ZnF2*^, *zld*^*JAZ*^.

### Cas9 genome engineering

#### Molecular reagents

*gRNAs* – gRNA sequences were identified using flyCRISPR Optimal Target Finder [42] and sequences are listed in Supplemental Table S3. Target-specific sequences for *zld* were cloned into the BbsI site of pU6-BbsIchiRNA [29].

*double-stranded DNA (dsDNA) donor* – The dsDNA donor templates for homologous recombination contained 1-kb homology arms flanking a *3xP3-DsRed* cassette [42] and a *sfGFP-* or *mCherry*- coding sequence downstream of the start codon of *zld-RB,* or a *mCherry* coding sequence just upstream of the stop codon of exon 2, which codes for the alternative splice isoform *zld-RD*.

*single-stranded donor oligonucleotide (ssODN)*– The ssODN donor templates for homologous recombination contained >50bp homology directly adjacent to the Cas9-mediated double stranded break and desired point mutations in the target locus. Silent mutations were engineered in the ssODN to mutate the gRNA target sequence and to introduce a novel restriction digest site. Integrated DNA Technologies synthesized the ssODNs.

Plasmids were purified with a HiSpeed Plasmid Midi Kit (Qiagen). Injection mixes were prepared in ultra-pure H2O and contained either 500ng/µl dsDNA donor plasmid and 250ng/µl gRNA plasmid, or 200ng/µl ssODN and 100ng/µl gRNA(s) plasmid. Constructs were injected into *w*^*1118*^; *PBac{y[+mDint2]=vas-Cas9}VK00027* (BDSC #51324) embryos. Injections were performed by Bestgene Inc.

#### Screening

*sfGFP-and mCherry-tagged zld isoforms – zld* alleles containing fluorescent tags were generated using Cas9-mediated genome editing and homology-directed repair from a dsDNA donor. Two gRNA plasmids and a dsDNA donor plasmid were injected into *vasa-Cas9* embryos (as described above). Engineered lines were identified by DsRed expression in the eye. piggyBac transposase was subsequently used to cleanly excise the DsRED marker [43] followed by sequence confirmation of precise tag incorporation.

*zld-RD deletion –* Two gRNA plasmids targeting sequence flanking the downstream coding exon for *the zld-RD* isoform and an ssODN containing attP sequence were injected into *vasa-Cas9* embryos (as described above). PCR was performed using primers flanking the target locus, and NHEJ products were resolved on a 0.8% agarose gel and visualized under UV light. *zld-RD* deletion alleles were confirmed by sequence analysis.

*zld targeted point mutations – *zld*^*ZnF1*^ zld*^*EDD*^, *zld*^*ZnF2*^, *zld*^*JAZ*^ – *zld* alleles containing point mutations within highly conserved domains were generated using Cas-9 mediated genome editing and homology-directed repair from a ssODN. A single gRNA plasmid and a ssODN were injected into *vasa-Cas9* embryos (as described above). Adults that developed from injected embryos were individually crosses to *w*^*1118*^. The offspring were crossed in batch (~10 flies per cross) to their siblings or FM7 balancer and given several days to mate. Genomic DNA was extracted from the batch of parental flies by homogenization in 300µl buffer (100mM Tris HCl pH7.5, 100mM EDTA pH8.0, 100mM NaCl, 0.5% SDS), followed by incubation at 65°C for 30 minutes. DNA was precipitated by adding 600µl 1:2.5 [5M]KOAc:[6M]LiCl, incubating on ice for 10 minutes, followed by centrifugation at 14000 x g for 15 minutes. Supernatant was collected, 450µl isopropanol was added and sample was centrifuged at 14000 x g for 15 minutes. DNA pellet was briefly washed with 70% ethanol and re-suspended in 36µl TE supplemented with 1mg/mL RNase A. Mutant alleles were screened for by PCR amplification followed by restriction digest. Digested bands corresponding to a mutant allele were resolved on an 8% TBE gel and stained with SYBR Gold nucleic acid gel stain (Invitrogen).

The offspring of batch crosses carrying the mutant allele were individually crossed to FM7 balancer and given several days to mate. Parental individuals were sacrificed, and DNA was extracted from single flies by homogenization in 50µl of buffer (10mM Tris-HCl pH 8., 1mM EDTA, 25mM NaCl, 200 μg/ml proteinase K), followed by incubation at 37°C for 30 minutes and inactivation of the proteinase K enzyme at 95°C for 2 minutes. Mutant alleles were screened for by PCR amplification followed by restriction digest. Digested bands were resolved on a 0.8% agarose gel and visualized under UV light. Transmission of expected genome editing events was confirmed by sequence analysis.

### *Drosophila* genetics

Complementation with previously identified recessive mutation *zld*^*294*^ [14] was performed to verify phenotypes were a result of Cas9-mediated mutagenesis within *zld*. Trans-heterozygous females were crossed to *w*^*1118*^ males to determine fertility.

*mat-α-GAL4-VP16* males were crossed with *UASp-zld* transgenic females. *w:mat-α-GAL4-VP16/+:UASp-zld/+* were recovered from previous crosses and mated to siblings. The percentage of embryos hatched was determined by lining up approximately two hundred and fifty 0-3 hour old embryos and counting the hatched eggs after at least 24 hours. All genetic experiments were carried out at 25°C.

### Confocal imaging

Embryos were dechorionated and analyzed under halocarbon oil to determine stage. Third instar larval (L3) wing discs were dissected and mounted in PBS. Confocal images were acquired on an A1R-S Confocal Microscope (Nikon) with 20x objective. Images were analyzed using NIS-Elements AR software. Z-stacks were flattened using the Maximum Intensity Z-projection function.

### RNA-sequencing and analysis

RNA-seq experiments were done on hand-sorted stage 5 embryos laid by *zld*^*ZnF2*^ (C554S mutation) homozygous females and *w*^*1118*^ embryos as a wild-type control. Embryos were dechorionated, analyzed under halocarbon oil to determine stage, collected and lysed in TRIzol (ThermoFisher) supplemented with 150 µg/ml glycogen. RNA was extracted, and cDNA libraries were prepared using Truseq RNA sample prep kit (Illumina). Three replicates of each were sequenced. The cDNA 100 bp single-end reads were sequenced at the UW Biotechnology Center DNA Sequencing Facility using an Illumina HiSeq 2000. Using the Galaxy platform [44], reads were examined for quality, trimmed, and filtered. The reads were then mapped to the BDGP D. melanogaster (dm6) genome using RNA-STAR (Galaxy Version 2.5.2b-0). Cufflinks (Galaxy Tool version 2.2.1) was used with default settings for transcript assembly. The resulting assembled transcripts were compared using Cuffdiff [45](R version 3.1.2) to identify genes that change significantly (p-value<0.05, >two-fold change) in expression. Only genes that were significantly mis-expressed in all replicates were used for further analysis.

Prior to comparisons, all gene IDs were converted to current FlyBase identifiers (FBgn#) using the ‘Upload/Convert IDs’ tool available on FlyBase [32]. Single-embryo expression data from Lott et al. 2010 [34] were used to classify up- or down-regulated genes in mutant embryos as (1) zygotic, (2) zygotic-maternal, and (3) maternal only. Translation and stability datasets of maternal mRNAs from Thomsen et al. 2010 [10] were used to classify up-regulated maternal genes as targets of maternal degradation or zygotic degradation. mRNAs degraded by the maternal pathway were considered as Classes II and III (‘exclusively maternally degraded’ and ‘maternally degraded and transcribed’); and mRNAs degraded by the zygotic pathway, were considered as Class IV (‘exclusively zygotically degraded’). ZLD ChIP-seq data from Harrison et al. 2011 [13] were used to identify the number of down-regulated zygotic genes bound by ZLD. Enrichments and depletions for comparisons to data from Lott et al. 2010 [34], Thomsen et al. 2010 [10], and Harrison et al. 2011 [13] were determined using a Fisher’s exact test.

### Gene ontology annotations

Gene ontology (GO) annotation was performed using the online GO Consortium tool (http://geneontology.org/), which uses the PANTHER classification system [46]. Lists of gene names were entered searching for enrichment in molecular function using a Bonferroni correction. The data were collected using the PANTHER over-representation test release 20170413 with the 2017-06-29 GO ontology database release.

## ACKNOWLEDGMENTS

The authors thank Kate O’Connor-Giles and Jill Wildonger for helpful discussions regarding Cas9-mediated genome editing, and Catherine Fox and Andy Mehle for comments on the manuscript. Thanks to Katharine Schulz for assistance in dissecting larval wing discs. We would also like to thank the University of Wisconsin Biotechnology Center DNA Sequencing Facility and the Biochemistry Optical Core for assistance with high-throughput sequencing and fluorescent imaging, respectively.

## References

1. Schier AF. The maternal-zygotic transition: death and birth of RNAs. Science. 2007;316(5823):406–7.

2. Tadros W, Lipshitz HD. The maternal-to-zygotic transition: a play in two acts. Development. 2009;136(18):3033–42.

3. Harrison MM, Eisen MB. Transcriptional Activation of the Zygotic Genome in Drosophila. Matern Zygotic Transit. 2015;113:85–112.

4. Lee MT, Bonneau AR, Giraldez AJ. Zygotic genome activation during the maternal-to-zygotic transition. Annu Rev Cell Dev Biol. 2014;30:581–613.

5. Bashirullah A, Halsell SR, Cooperstock RL, Kloc M, Karaiskakis A, Fisher WW, et al. Joint action of two RNA degradation pathways controls the timing of maternal transcript elimination at the midblastula transition in Drosophila melanogaster. EMBO J. 1999 May 4;18(9):2610–20.

6. Tadros W, Houston SA, Bashirullah A, Cooperstock RL, Semotok JL, Reed BH, et al. Regulation of maternal transcript destabilization during egg activation in Drosophila. Genetics. 2003 Jul;164(3):989–1001.

7. Aanes H, Winata CL, Lin CH, Chen JP, Srinivasan KG, Lee SGP, et al. Zebrafish mRNA sequencing deciphers novelties in transcriptome dynamics during maternal to zygotic transition. Genome Res. 2011 Aug;21(8):1328–38.

8. Audic Y, Anderson C, Bhatty R, Hartley RS. Zygotic Regulation of Maternal Cyclin A1 and B2 mRNAs. Mol Cell Biol. 2001 Mar 1;21(5):1662–71.

9. Hamatani T, Carter MG, Sharov AA, Ko MSH. Dynamics of Global Gene Expression Changes during Mouse Preimplantation Development. Dev Cell. 2004 Jan;6(1):117–31.

10. Thomsen S, Anders S, Janga SC, Huber W, Alonso CR. Genome-wide analysis of mRNA decay patterns during early Drosophila development. Genome Biol. 2010;11(9):R93.

11. Yartseva V, Giraldez AJ. The Maternal-to-Zygotic Transition During Vertebrate Development: A Model for Reprogramming. Curr Top Dev Biol. 2015;113:191–232.

12. Laver JD, Li X, Ray D, Cook KB, Hahn NA, Nabeel-Shah S, et al. Brain tumor is a sequence-specific RNA-binding protein that directs maternal mRNA clearance during the Drosophila maternal-to-zygotic transition. Genome Biol. 2015 May 12;16(1):94.

13. Harrison MM, Li X-Y, Kaplan T, Botchan MR, Eisen MB. Zelda Binding in the Early Drosophila melanogaster Embryo Marks Regions Subsequently Activated at the Maternal-to-Zygotic Transition. Copenhaver GP, editor. PLoS Genet. 2011 Oct 20;7(10):e1002266.

14. Liang HL, Nien CY, Liu HY, Metzstein MM, Kirov N, Rushlow C. The zinc-finger protein Zelda is a key activator of the early zygotic genome in Drosophila. Nature. 2008;456(7220):400–3.

15. Nien C-Y, Liang H-L, Butcher S, Sun Y, Fu S, Gocha T, et al. Temporal coordination of gene networks by Zelda in the early Drosophila embryo. Barsh GS, editor. PLoS Genet. 2011 Oct 20;7(10):e1002339.

16. Fu S, Nien C-Y, Liang H-L, Rushlow C. Co-activation of microRNAs by Zelda is essential for early Drosophila development. Development. 2014;141(10):2108–18.

17. Staudt N, Fellert S, Chung H-R, Jäckle H, Vorbrü G. Mutations of the Drosophila Zinc Finger– encoding Gene vielfä ltig Impair Mitotic Cell Divisions and Cause Improper Chromosome Segregation □ V. Mol Biol Cell. 2006;17:2356–65.

18. Eichhorn SW, Subtelny AO, Kronja I, Kwasnieski JC, Orr-Weaver TL, Bartel DP. mRNA poly(A)-tail changes specified by deadenylation broadly reshape translation in Drosophila oocytes and early embryos. Elife. 2016 Jul 30;5.

19. Bushati N, Stark A, Brennecke J, Cohen SM. Temporal reciprocity of miRNAs and their targets during the maternal-to-zygotic transition in Drosophila. Curr Biol. 2008 Apr 8;18(7):501–6.

20. Hamm DC, Bondra ER, Harrison MM. Transcriptional Activation Is a Conserved Feature of the Early Embryonic Factor Zelda That Requires a Cluster of Four Zinc Fingers for DNA Binding and a Low-complexity Activation Domain. J Biol Chem. 2015 Feb 6;290(6):3508–18.

21. ten Bosch JR, Benavides JA, Cline TW. The TAGteam DNA motif controls the timing of Drosophila pre-blastoderm transcription. Development. 2006 May 12;133(10):1967–77.

22. Bradley RK, Li X-Y, Trapnell C, Davidson S, Pachter L, Chu HC, et al. Binding site turnover produces pervasive quantitative changes in transcription factor binding between closely related Drosophila species. Wray GA, editor. PLoS Biol. 2010 Mar 23;8(3):e1000343.

23. Biedler JK, Hu W, Tae H, Tu Z. Identification of Early Zygotic Genes in the Yellow Fever Mosquito Aedes aegypti and Discovery of a Motif Involved in Early Zygotic Genome Activation. Aerts S, editor. PLoS One. 2012 Mar 23;7(3):e33933.

24. Paris M, Kaplan T, Li XY, Villalta JE, Lott SE, Eisen MB. Extensive divergence of transcription factor binding in Drosophila embryos with highly conserved gene expression. Wittkopp P, editor. PLoS Genet. 2013 Sep 12;9(9):e1003748.

25. Ribeiro L, Tobias-Santos V, Santos D, Antunes F, Feltran G, de Souza Menezes J, et al. Evolution and multiple roles of the Pancrustacea specific transcription factor zelda in insects. Desplan C, editor. PLOS Genet. 2017 Jul 3;13(7):e1006868.

26. Roy S, Ernst J, Kharchenko P V., Kheradpour P, Negre N, Eaton ML, et al. Identification of Functional Elements and Regulatory Circuits by Drosophila modENCODE. Science (80-). 2010 Dec 24;330(6012):1787–97.

27. Pearson JC, Watson JD, Crews ST. Drosophila melanogaster Zelda and Single-minded collaborate to regulate an evolutionarily dynamic CNS midline cell enhancer. Dev Biol. 2012 Jun 15;366(2):420–32.

28. Giannios P, Tsitilou SG. The embryonic transcription factor Zelda of Drosophila melanogaster is also expressed in larvae and may regulate developmentally important genes. Biochem Biophys Res Commun. 2013 Aug 23;438(2):329–33.

29. Gratz SJ, Cummings AM, Nguyen JN, Hamm DC, Donohue LK, Harrison MM, et al. Genome engineering of Drosophila with the CRISPR RNA-guided Cas9 nuclease. Genetics. 2013;194(4):1029–35.

30. Fisher B, Weiszmann R, Frise E, Hammonds A, Tomancak P, Beaton A, et al. BDGP insitu homepage. 2012.

31. Struffi P, Corado M, Kaplan L, Yu D, Rushlow C, Small S. Combinatorial activation and concentration-dependent repression of the Drosophila even skipped stripe 3+7 enhancer. Development. 2011 Oct 1;138(19):4291–9.

32. Gramates LS, Marygold SJ, Santos G Dos, Urbano J-M, Antonazzo G, Matthews BB, et al. FlyBase at 25: looking to the future. Nucleic Acids Res. 2017 Jan 4;45(D1):D663–71.

33. Yang M, May WS, Ito T. JAZ requires the double-stranded RNA-binding zinc finger motifs for nuclear localization. J Biol Chem. 1999 Sep 24;274(39):27399–406.

34. Lott SE, Villalta JE, Schroth GP, Luo S, Tonkin LA, Eisen MB. Noncanonical Compensation of Zygotic X Transcription in Early Drosophila melanogaster Development Revealed through Single-Embryo RNA-Seq. Hawley RS, editor. PLoS Biol. 2011 Feb 8;9(2):e1000590.

35. Schulz KN, Bondra ER, Moshe A, Villalta JE, Lieb JD, Kaplan T, et al. Zelda is differentially required for chromatin accessibility, transcription-factor binding and gene expression in the early Drosophila embryo. Genome Res. 2015;1715–26.

36. Arsala D, Lynch JA. Ploidy has little effect on timing early embryonic events in the haplodiploid wasp Nasonia. genesis. 2017 May;55(5):e23029.

37. Miura T, Satoh T, Takeuchi H. Role of metal-ligand coordination in the folding pathway of zinc finger peptides. Biochim Biophys Acta. 1998 Apr 23;1384(1):171–9.

38. Burge RG, Martinez-Yamout MA, Dyson HJ, Wright PE. Structural Characterization of Interactions between the Double-Stranded RNA-Binding Zinc Finger Protein JAZ and Nucleic Acids. Biochemistry. 2014 Mar 11;53(9):1495–510.

39. Schneider I. Cell lines derived from late embryonic stages of Drosophila melanogaster. J Embryol Exp Morphol. 1972 Apr;27(2):353–65.

40. Harrison MM, Botchan MR, Cline TW. Grainyhead and Zelda compete for binding to the promoters of the earliest-expressed Drosophila genes. Dev Biol. 2010;345(2):248–55.

41. Wright KJ, Marr MT, Tjian R. TAF4 nucleates a core subcomplex of TFIID and mediates activated transcription from a TATA-less promoter. Proc Natl Acad Sci. 2006 Aug 15;103(33):12347–52.

42. Gratz SJ, Ukken FP, Rubinstein CD, Thiede G, Donohue LK, Cummings AM, et al. Highly Specific and Efficient CRISPR/Cas9-Catalyzed Homology-Directed Repair in Drosophila. Genetics. 2014 Apr;196(4):961–71.

43. Bruckner JJ, Zhan H, Gratz SJ, Rao M, Ukken F, Zilberg G, et al. Fife organizes synaptic vesicles and calcium channels for high-probability neurotransmitter release. J Cell Biol. 2017 Jan 2;216(1):231–46.

44. Afgan E, Baker D, van den Beek M, Blankenberg D, Bouvier D, Čech M, et al. The Galaxy platform for accessible, reproducible and collaborative biomedical analyses: 2016 update. Nucleic Acids Res. 2016 Jul 8;44(W1):W3–10.

45. Trapnell C, Hendrickson DG, Sauvageau M, Goff L, Rinn JL, Pachter L. Differential analysis of gene regulation at transcript resolution with RNA-seq. Nat Biotechnol. 2012 Dec 9;31(1):46–53.

46. Ashburner M, Ball CA, Blake JA, Botstein D, Butler H, Cherry JM, et al. Gene ontology: tool for the unification of biology. The Gene Ontology Consortium. Nat Genet. 2000 May;25(1):25–9.

